# Deep Brain Stimulation for Parkinson’s disease changes perception in the Rubber Hand Illusion

**DOI:** 10.1101/231340

**Authors:** Catherine Ding, Colin J Palmer, Jakob Hohwy, George J Youssef, Bryan Paton, Naotsugu Tsuchiya, Julie C Stout, Dominic Thyagarajan

## Abstract

Parkinson’s disease (PD) alters cortico-basal ganglia-thalamic circuitry and susceptibility to an illusion of bodily awareness, the Rubber Hand Illusion (RHI). Bodily awareness is thought to result from multisensory integration in a predominantly cortical network; the role of subcortical connections is unknown. We studied the effect of modulating cortico-subcortical circuitry on multisensory integration for bodily awareness in PD patients treated with subthalamic nucleus (STN) deep brain stimulation (DBS) using the RHI experiment. Typically, synchronous visuo-tactile cues induce a false perception of touch on the rubber hand as if it were the subject’s hand, whereas asynchronous visuo-tactile cues do not. However, we found that in the asynchronous condition, patients in the off-stimulation state did not reject the RHI as strongly as healthy controls; switching on STN-DBS partially ‘normalised’ their responses. Patients in the off-stimulation state also misjudged the position of their hand, indicating it to be closer to the rubber hand than controls. However, STN-DBS did not affect proprioceptive judgements or subsequent arm movements altered by the perceptual effects of the illusion. Our findings support the idea that the STN and subcortical connections have a key role in multisensory integration for bodily awareness. Decisionmaking in multisensory bodily illusions is discussed.

## Introduction

Bodily awareness is thought to result from the integration of multiple sensory cues with an internal sense of time and body image.^1^ Prefrontal, premotor and intraparietal cortices and the cerebellum have been implicated in a neural network for multisensory integration in body awareness.^2–4^ However, little is known about the subcortical structures that may underpin this network.

We suspect that subcortical structures may be involved in multisensory integration for bodily awareness because people with Parkinson’s disease (PD) have an atypical reaction to the Rubber Hand Illusion (RHI).^5^ In the RHI paradigm, synchronous visual and tactile cues (from stroking the subject’s hidden hand and a visible rubber hand) typically induce a false perception of touch on the rubber hand as if it were the subject’s hand, whereas asynchronous visual and tactile cues do not elicit this illusion. Also, subjects mislocalize their unseen hand closer to the position of the rubber hand in the synchronous condition compared with the asynchronous condition (measured as “proprioceptive drift”). Previously, we reported that people with PD have a similar RHI experience to healthy controls in the synchronous condition, but fail to reject the RHI experience as strongly in the asynchronous condition.^5^ PD patients also have increased proprioceptive drift towards the rubber hand independent of visual-tactile synchrony.^5^ Together these findings suggest that PD increases susceptibility to altered bodily awareness. The mechanism for this unusual RHI response is unknown.^1^

The pathophysiological basis of PD is striatal dopaminergic degeneration resulting in altered cortico-basal ganglia-thalamic circuitry.^6^ Hence, PD is treated with dopaminergic drugs and high-frequency deep brain stimulation (DBS) of the subthalamic nucleus (STN) or globus pallidus interna (GPi), which is thought to work by modulating cortico-basal ganglia-thalamic circuitry.^7^ Previously, we reported that dopaminergic drugs did not affect either the subjective RHI or proprioceptive drift.^5^ This study explores the possible subcortical mechanisms of bodily awareness by examining the effect of STN-DBS on the RHI.

The effects of STN-DBS on sensory perception are not well understood, but there are suggestions that stimulation may affect multisensory integration for bodily awareness. Firstly, STN-DBS improves accuracy in somatosensory tasks in PD: switching on stimulation improves tactile detection thresholds^8^, proprioceptive accuracy^9^ and the integration of proprioceptive and tactile cues in haptic perception.^10^ Secondly, STN-DBS improves temporal perception: switching on stimulation improves temporal estimation of auditory intervals.^11,12^

However, PD patients treated with STN-DBS demonstrate subtle cognitive deficits in comparison to PD patients who have not undergone STN-DBS,^13^ suggesting that STN-DBS may affect higher-order perception. In particular, STN-DBS accelerates perceptual and cognitive decisions in high-conflict situations, which can result in more errors.^14–16^ Will a tendency to “jump to conclusions” increase or reduce illusory ownership when patients with STN-DBS process the complex and conflicting visual-tactile-proprioceptive cues of the RHI?

To answer this question and to probe the role of the STN and cortico-basal ganglia-thalamic circuitry on multisensory integration for body awareness, we studied the RHI in PD patients treated with STN-DBS. Based on our previous findings in PD patients without STN-DBS,^5^ we expected STN-DBS treated patients would have: 1) higher illusion scores on the RHI questionnaire than controls in the asynchronous condition, with no differences in the synchronous condition, and 2) greater proprioceptive drift than controls in both stroking conditions.

Our first hypothesis was that when STN-DBS is switched on, patients will have lower illusion scores in the asynchronous condition compared with when STN-DBS is switched off. We reasoned that improved processing of visual, somatosensory and temporal cues on Stimulation^8–11,17–19^would lead to a more typical rejection of the RHI when visual-tactile cues are asynchronous. We expected to see this only in the asynchronous condition because, in our previous study, PD had no effect on illusion scores in the synchronous condition.^5^

In our previous study^5^, proprioceptive drift increased in both synchronous and asynchronous conditions. In a study of kinesthesia in PD, STN-DBS improved proprioceptive deficits.^9^ Thus, our second hypothesis was that switching STN-DBS on would decrease proprioceptive drift in both stroking conditions. To differentiate the effect of STN-DBS implantation surgery on the RHI from that of STN stimulation itself, we tested eligible subjects before and after they underwent implantation surgery.

Lastly, we included a reaching task at the end of each RHI trial to assess the influence of somatic illusions on movement. For healthy subjects, the literature is mixed on whether the RHI affects the trajectory of reaching tasks^20–22^ or pointing accuracy.^4,23^, We had expected the RHI to have a greater influence on the subsequent actions of PD patients than controls because patients have proprioceptive deficits that compromise reaching accuracy^24^; we found no such effect in either patients or controls in our previous study^5^. However, most of those patients were mildly affected. Here, we examined the effect of the RHI on subsequent movement in STN-DBS-treated PD patients, who not only have more advanced disease, but in whom we can manipulate motor function by switching stimulation on or off. Thus, our third hypothesis was that the reach-to-grasp performance of STN-DBS-treated patients would differ between the synchronous and asynchronous condition, in addition to the expected improvement in motor function from stimulation of the STN.^25^

Investigating these hypotheses will inform the role of the STN and subcortical structures in multisensory integration for body awareness and the decision to reject illusory perceptions of self. A practical benefit of our study is that it may help clinicians understand the perceptual consequences of DBS, an increasingly utilised therapy for PD.

## Methods

### Participants

We recruited 24 patients with idiopathic PD treated with STN-DBS from our Movement Disorders Clinic, excluding those with clinically significant sensory or cognitive deficits. 12 of these patients had only postoperative testing. The other 12 patients had STN-DBS implantation surgery within the time frame of this study and were opportunistically tested pre- and postoperatively. Preoperative data from eight of these 12 patients and data from 21 healthy controls (aged 50-80) in this study has been published as part of a previous study.^5^ Participant characteristics are shown in Table 1. Compared with the PD patients in this study, controls were older and showed less symptomatology on the Hospital Anxiety and Depression Scale (HADS) and the Apathy Scale, but Montreal Cognitive Assessment (MoCA) scores were similar. This study was approved by the Monash Health Research Ethics Committee (HREC 12350B) and carried out in accordance with national ethical guidelines. All subjects gave written informed consent.

**Table 1.** Demographic and clinical characteristics of all 45 subjects (see text for details). UPDRS: Movement Disorders Society Unified Parkinson’s Disease Rating Scale, SD: standard deviation, IQR: interquartile range. * Parkinson’s disease patients were tested in the more severely affected hand. In controls, the hand tested was matched to patients. ** Four patients were not taking any dopaminergic drugs postoperatively.

|  | Parkinson's disease patients with STN-DBS | Controls | Test Statistic | *p* value |
| --- | --- | --- | --- | --- |
| Number | 24 | 21 |  |  |
| Sex (male) | 15 | 10 | $\chi^2 = 1.00$ | .32 |
| Handedness (right) | 17 | 19 | $\chi^2 = 2.70$ | .10 |
| Median Age (IQR) | 61 (56-63.75) | 66 (61-70) | Wilcoxon rank sum $W = 349$ | .03 |
| Median Montreal Cognitive Assessment score (IQR) | 28 (27-28.5) | 27 (26-28) | Wilcoxon rank sum $W = 190.5$ | .23 |
| Median Hospital Anxiety and Depression Scale score (IQR) | 9.5 (7.25-15) | 7 (3 - 10) | Wilcoxon rank sum $W = 146$ | .04 |
| Median Apathy Scale score (IQR) | 15.5 (7.5-18) | 7 (4-10) | Wilcoxon rank sum $W = 108$ | .01 |
| Median Parkinson's disease duration in years (IQR) | 13.5 (10-20.25) | NA |  |  |
| Hoehn and Yahr Parkinson's disease stage | 2-4 | NA |  |  |
| Side-tested (right hand)* | 11 | 10 |  |  |
| Median STN-DBS duration in months (IQR) | 16.5 (7-38) | NA |  |  |
| Median levodopa equivalent daily dose in mg(IQR)** | 416.5 (200-599.2) | NA |  |  |
| Median UPDRS Part III 'off' (IQR) | 35 (27.5-42.75) | NA | Wilcoxon signed rank of 'on' vs. 'off' scores $V = 0$ | <.0001 |
| Median UPDRS Part III 'on' (IQR) | 16 (8.5-21) | NA |  |  |

For the entire patient cohort, the median disease duration from diagnosis was 13.5 years (IQR 10-20 years). The median duration of STN-DBS treatment was 16.5 months (IQR 7-38 months) with a minimum of 5 months. Postoperatively, all patients had a CT brain scan within 24 hours, which was then fused with their preoperative MRI brain scan to confirm lead placement within the STN (Fig. 1A). A neurologist (CD) assessed motor severity (on- and off-stimulation) using the Movement Disorders Society Unified Parkinson’s Disease Rating Scale Part III (UPDRS).

**Figure 1.**
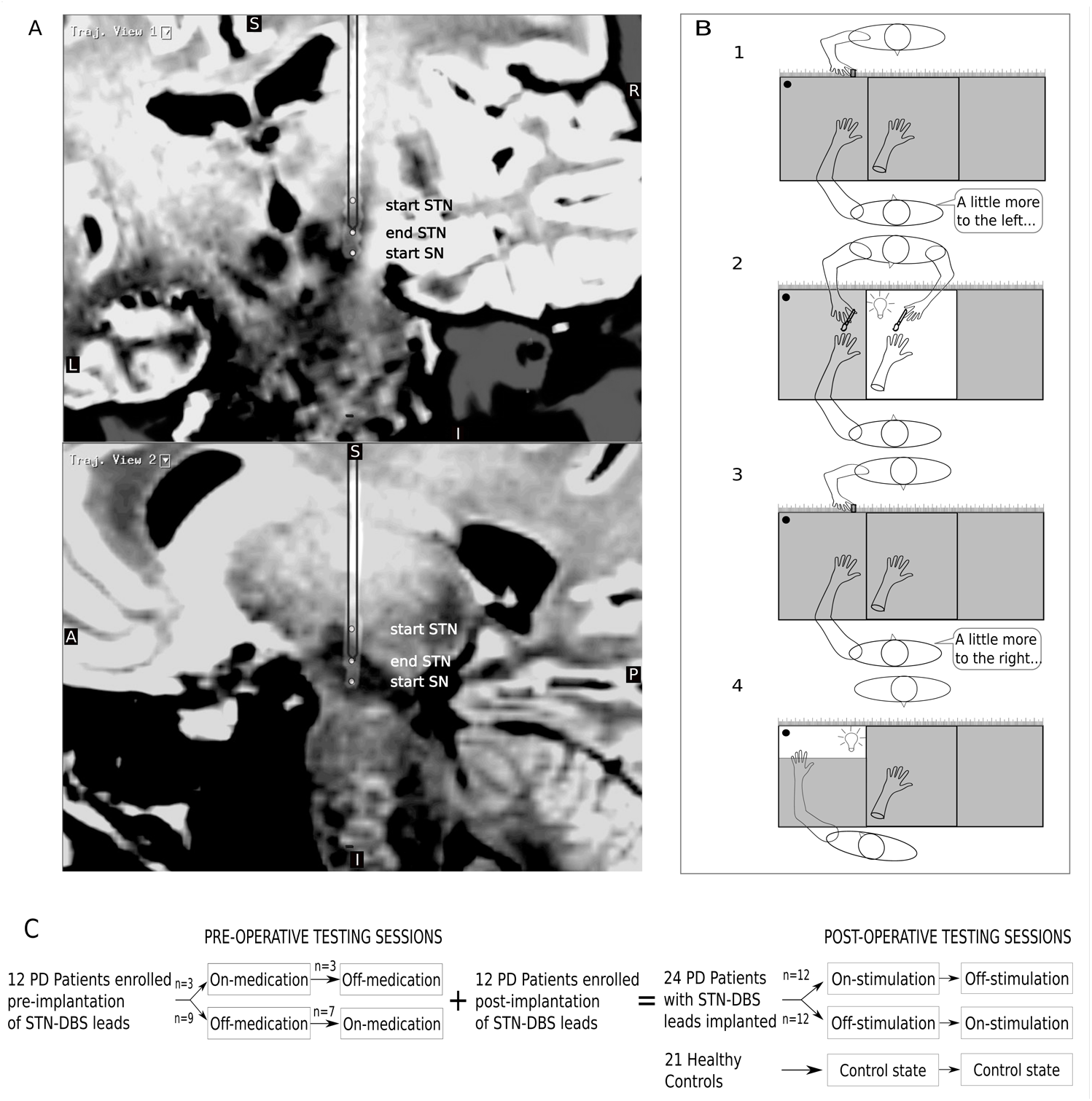
A) Example postoperative CT fused with the preoperative MRI (sagittal 3D FLAIR sequences with 1mm slices obtained on a Siemens Verio 3 Tesla unit). Orthogonal planes intersecting through a right Medtronics™ 3387 lead are shown (S: superior; I: inferior; L: left; R: right; A: anterior; P: posterior). White dots map the subthalamic nucleus (STN) and substantia nigra (SN) according to intraoperative semi-microelectrode recordings. B) Graphical timeline of a trial: 1. Subject verbally indicates their pre-stroking estimate of middle finger position, 2. Investigator either synchronously or asynchronously (randomised) strokes the subject’s hand (shaded = hidden) and a visible rubber hand, 3. Subject verbally indicates their post-stroking estimate of middle finger position, 4. Subject reaches for a cylinder now illuminated in the far corner of the lateral compartment. At the end of each trial, the subject fills in the Rubber Hand Illusion questionnaire (not pictured). C) Graphical timeline of the study. Pre-DBS implantation surgery, patients were tested in on and off-medication states. Post-DBS implantation surgery, patients were tested in on and off-stimulation states. Controls were also tested twice. Fig. 1B is from Ding, C. et al. (2017) Parkinson’s disease alters multisensory perception: Insights from the Rubber Hand Illusion, Neuropsychologia by Elsevier Science Ltd. Reproduced with permission of Elsevier Science Ltd in the format Journal/magazine via Copyright Clearance Center.

In the 12 subjects with pre-operative testing, the median interval between pre- and postoperative testing was 12.5 months (IQR 9.75-17.25 months). Although subjects took less dopaminergic medication postoperatively (preoperative levodopa equivalent daily dose (LEDD): Mdn (IQR) = 1350 (1090-1973); postoperative LEDD: Mdn (IQR) = 417 (0-508), *V* = 75, *p* = .005), there was no difference in motor severity in respective on-treatment states (preoperative on-medication UPDRS: Mdn (IQR) = 15 (11.5-16); postoperative on-stimulation UPDRS: Mdn (IQR) = 14 (9.25-16.25), *V* = 22, *p* = 1) or off-treatment states (preoperative off-medication UPDRS: Mdn (IQR) = 31.5 (18-28); postoperative off-stimulation UPDRS: Mdn (IQR) = 33 (22.5-41), *V* = 25.5, *p* = .878). Implantation surgery did not significantly affect scores on the MoCA, HADS or Apathy Scale (Supplementary Table A)

### Experiment Design

The RHI set-up used in this study is detailed in our previous paper^5^ and the graphical timeline of a trial is reproduced in Fig. 1B. Briefly, subjects sat in front of a black box with three compartments and a dark glass top that allowed them to view the contents of each compartment only when individually illuminated. In each trial, the hand being tested was placed in the corresponding lateral compartment, while an identically aligned rubber hand was placed in the central compartment. Each trial began with the subject making a ‘baseline estimate’ of the position of their unseen middle finger. They did this by verbally directing the investigator to move a marker along the length of the box until they were satisfied that the marker was directly in front of their middle finger. Then, the central compartment was illuminated to reveal the rubber hand. The subject was instructed to watch the rubber hand as the investigator stroked it alongside their hidden hand. In the synchronous condition, the experimenter stroked corresponding fingers on the real and rubber hands in unison. In the asynchronous condition, the experimenter alternately stroked non-corresponding fingers on the real and rubber hands (i.e., such that there was both temporal and spatial asynchrony between the seen and felt touch). After two minutes of stroking, the light was switched off to hide the rubber hand once more. The subject then made a verbal ‘post-stroking estimate’ of the position of their hidden middle finger. Next, a cylinder in the far corner of the lateral compartment was illuminated, and the subject was instructed to touch the cylinder. Their hand was hidden from view until it neared the cylinder. At the end of each trial, the subject took their hand out of the box and completed a RHI questionnaire that assessed their perceptual experiences during that trial.

A testing session comprised two trials of the synchronous condition and two trials of the asynchronous condition in randomised order. Patients were tested on the more severely affected hand; in controls the hand tested was matched to patients, ensuring each group had the same number of left and right-sided trials. All subjects completed at least two testing sessions (Fig. 1C). For STN-DBS patients, one session commenced after stimulation had been switched off for 30 min (‘off-stimulation’) while the other session commenced after stimulation had been switched on at usual settings for at least 30 min. In a counterbalanced, crossover design, half the STN-DBS patients were tested on-stimulation first, while the other half were tested in the reverse order. Patients took their usual anti-Parkinson medication in the on- and off-stimulation states. The subset of 12 patients studied preoperatively were tested after their dopaminergic drugs were withheld for at least 12 hours (‘off-medication’). In 10 of these patients, we had the opportunity to complete another preoperative testing session while they were taking their usual dopaminergic drugs (‘on-medication’). This enabled us to make a comprehensive comparison of pre- and postoperative on and off-states achieved by medication and STN-DBS respectively.

### Measures

We recorded three outcome measures from each trial: (1) questionnaire responses (2) proprioceptive drift, and (3) reach movement metrics.

### Questionnaire responses

Subjects completed an 11 statement RHI questionnaire used in previous studies,^5,22,26^ comprising three critical (‘illusion’) statements designed to elicit the subjective strength of the RHI, intermixed with eight control (‘mock’) statements designed to detect response biases (Supplementary Table B). Subjects marked their response on a 14 cm long visual analogue scale with “Strongly disagree” at 0 cm, “Strongly agree” at 14 cm and “Very unsure whether to agree or disagree” in the middle. Thus, higher illusion scores quantitatively capture endorsement of the RHI, while lower illusion scores quantify rejection of the RHI.

### Proprioceptive drift

We measured proprioceptive drift as the difference between the subject’s ‘baseline’ and ‘post-stroking’ estimates of middle finger position. We also assessed proprioceptive (in)accuracy as the difference between the ‘baseline’ estimate and the actual position of their middle finger. For both proprioceptive drift and baseline proprioceptive inaccuracy, positive measurements indicate the medial direction (towards the position of the rubber hand), whereas negative measurements indicate the lateral direction.

### Reach Movement Metrics

We recorded reach trajectories using a magnetic 3D motion tracker and a sensor attached to the subject’s thumbnail. We defined the reaching movement as the period starting from when the velocity of the sensor first exceeded 20 mm/s, to when it finally slowed below 20 mm/s. We then calculated seven reach metrics: 1. duration of movement, 2. mean velocity of movement, 3. peak velocity of movement, 4. time to reach peak velocity, 5. peak lateral displacement, 6. initial angle of movement (the angle of instantaneous velocity from the midline when 10% of the movement was completed) and 7. integrated jerk (the time derivative of acceleration integrated over the course of the movement).

### Statistical Analysis

To compare STN-DBS patients in ‘on-stimulation’ and ‘off-stimulation’ states with controls, we estimated mixed-effects regression models for questionnaire responses and proprioceptive drift using fixed effects for ‘subject group’ (controls vs. PD on-stimulation vs. PD off-stimulation), ‘stroking condition’ (synchronous vs. asynchronous) and their interaction. We estimated random effects for ‘stroking condition’ nested within subjects. Questionnaire data (visual analogue scores) was bound between 0 and 14 and thus treated as the odds of a binary response between strongly disagree and strongly agree. We used logistic regression on averaged responses to mock and illusion statements in each trial, estimating additional fixed and random effects for the statement type (‘mock’ vs. ‘illusion’). Linear regression was appropriate for proprioceptive drift data. We applied Bonferroni corrections for multiple comparisons to posthoc contrasts. We controlled for age, sex, cognition (MoCA), mood (HADS) and apathy (Apathy Scale) as potential covariates in both models. We also controlled for baseline proprioceptive inaccuracy in the proprioceptive drift model.

To evaluate the effect of STN-DBS implantation surgery on the RHI, in the subset of patients who had preoperative testing, we compared their outcome measures in four ‘treatment states’ (pre-op off-medication, pre-op on-medication, post-op off-stimulation, post-op on-stimulation). We estimated additional mixed effects models of questionnaire responses and proprioceptive drift, using the fixed effects of ‘treatment state’, ‘stroking condition’ and their interaction. Random effects for questionnaire response and proprioceptive drift models were as described above. We controlled for the duration of disease, motor severity (UPDRS) and LEDD in both models, as well as baseline proprioceptive inaccuracy in the proprioceptive drift model.

To evaluate the effect of the RHI on movement in the STN-DBS patients, we estimated linear mixed effects models for each of the seven reach movement metrics, using fixed effects of side-of-test, ‘stroking condition’, ‘stimulation state’ (on vs. off), and their interactions. Random effects were estimated for stimulation state nested within subjects. We controlled for motor severity (UPDRS) as a potential covariate. Statistical analysis was performed in R,^27^ using packages ‘lme4’,^28^ ‘nlme’,^29^ ‘Effects’,^30,31^ ‘Phia’,^32^ and ‘Stargazer’.^33^

The datasets generated during and/or analysed during the current study are available in the figshare repository, https://doi.org/10.6084/m9.figshare.5662219.v1.

## Results

### Subject excluded

One PD patient (who was studied preoperatively) had a unique postoperative RHI response wherein he strongly endorsed illusion statements in the asynchronous condition but not in the synchronous condition. When questioned at the conclusion of testing, he stated that in the synchronous trials he was watching his hand being stroked rather than a rubber hand! This response was not seen in any other subjects in our study, and to our knowledge, has not been reported in the RHI literature. Because of doubt over the validity of his questionnaire responses, we excluded him from all analysis.

### Questionnaire data

#### All subjects

As expected, all subject groups reported a stronger RHI in the synchronous condition compared with the asynchronous condition, which was reflected in the illusion statements but not the mock statements. Regression analysis of questionnaire responses showed a three-way interaction between ‘statement type’, ‘stroking condition’ and ‘subject group’ (χ2 = 37.35, *p* < .0001). Examination of contrasts revealed that all subject groups were more likely to agree with illusion statements in the synchronous condition than in the asynchronous condition (controls: χ2 = 156.67, *p* < .0001; PD off-stimulation: χ2 = 33.49, *p* < .0001; PD on-stimulation: χ2 = 68.38, *p* < .0001). By contrast, agreement with mock statements did not differ between stroking conditions or amongst subject groups (all *p* > .05). All regression tables are presented in Supplementary Material.

Focusing on the illusion statements (Fig 2A), as predicted from our previous study, PD patients did not reject the RHI in the asynchronous condition as strongly as controls (PD offstimulation: χ2 = 34.18, *p* < .0001; PD on-stimulation: χ2 = 12.26, *p* < .006). In the asynchronous condition, patients had lower illusion scores in the on-stimulation state compared with the off-stimulation state (χ2 = 20.02, *p* < .0001), which is consistent with our first hypothesis that switching on STN-DBS strengthens rejection of the RHI experience in the asynchonous condition. By contrast, in the synchronous condition, endorsement of the RHI was similar between controls and patient on- or off-stimulation (all *p* > .05). Age, sex, cognition, mood and apathy were not significant covariates and thus excluded from the final model (Supplementary Table C1-2).

**Figure 2.**
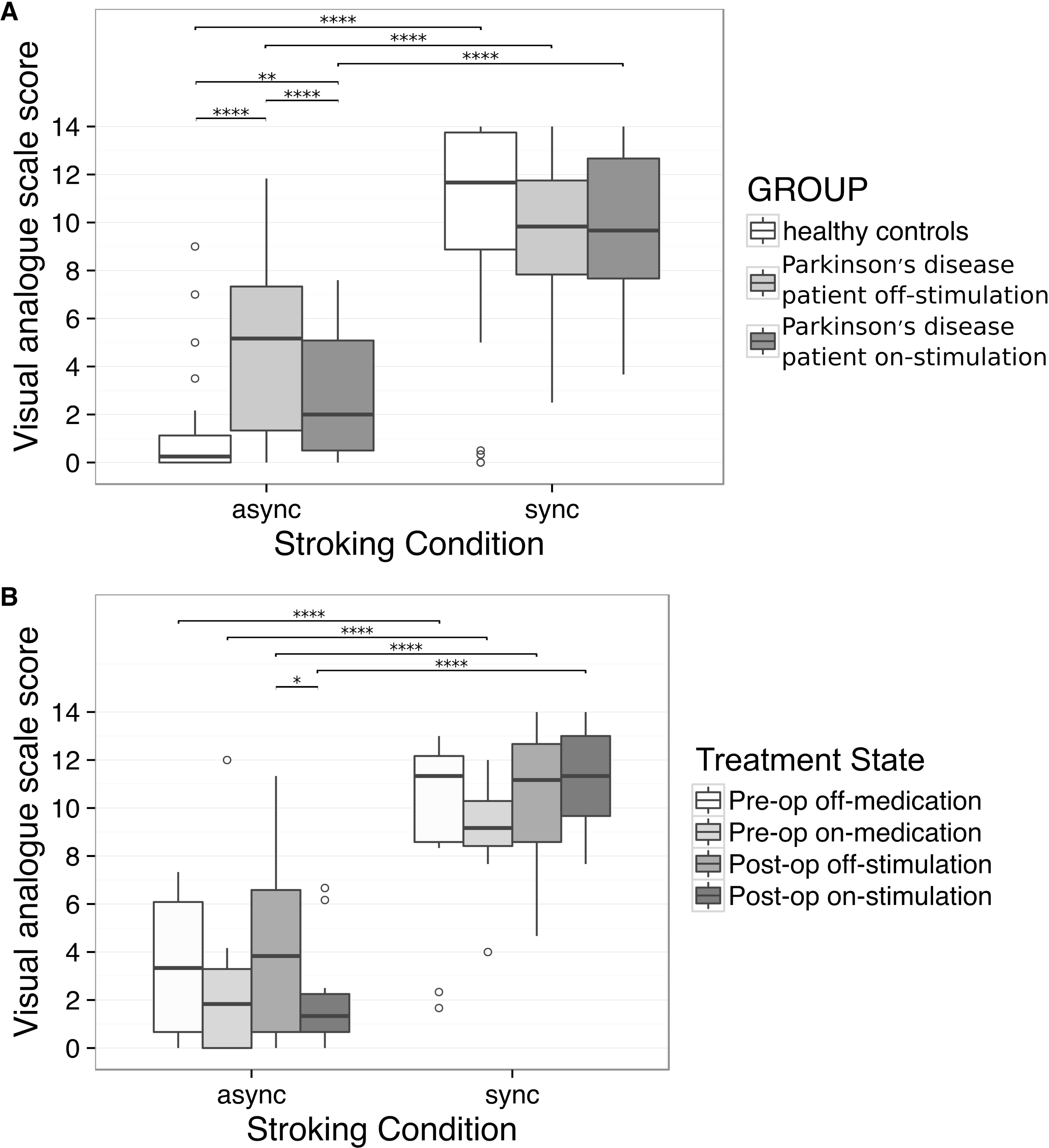
Scoring of critical Rubber Hand Illusion statements in A) 23 Parkinson’s disease patients treated with deep brain stimulation of the subthalamic nucleus (in on-stimulation and off-stimulation states) and 21 healthy controls and B) a subset of 11 Parkinson’s disease patients studied prior to STN-DBS implantation surgery (pre-op) whilst taking their usual anti-Parkinson drugs (‘on-medication’, n = 9), and after withholding all anti-Parkinson drugs (‘off-medication’, n = 11). Thick line: median, hinge: IQR, whisker: hinge ± 1.5*IQR, °: outlier, async: asynchronous, sync: synchronous. P-values are based on the regression model, ****: p < .0001, **: p < .01, *p < .05. All unlabelled contrasts were not significant.

#### Subset of PD patients studied preoperatively

The preoperative subset of 11 patients had similar results to the larger patient cohort; regression analysis of questionnaire responses showed a three-way interaction between ‘statement type’, ‘stroking condition’ and ‘treatment state’ (χ2 = 19.06, *p* = .025). In all treatment states, patients had higher illusion scores in the synchronous stroking condition than in the asynchronous condition (pre-op off-medication: χ2 = 36.21, *p* < .0001; pre-op on-medication: χ2 = 32.83, *p* < .0001; post-op off-stimulation: χ2 = 36.17, *p* < .0001); post-op on-stimulation: χ2 = 77.44, *p* < .0001). By contrast, agreement with mock statements did not differ between stroking conditions in any treatment state (all *p* >.05).

Focussing on illusion statements (Fig. 2B), illusion scores did not differ between pre-op and post-op treatment states (all *p* > .05). As with the 23 STN-DBS patients, postoperative illusion scores in the asynchronous condition in this subset were lower when their stimulators were switched on (χ2 = 10.89, *p* = .023). Preoperative illusion scores in the asynchronous condition were similar between on- and off-medication states (*p* > .05). Illusion scores in the synchronous condition were similar between all four treatment states (all *p* > .05). LEDD was not a significant covariate and although duration of disease was significant (log(OR) = 0.07, 95% CI [0.02,0.13], *p* = .004), it did not affect the fixed effects of interest (Supplementary Table D1-2).

### Proprioceptive drift

#### All subjects

Replicating our previous findings,^5^ proprioceptive drift was larger in the synchronous condition than in the asynchronous condition, and in PD patients compared with controls (Fig. 3A). However, in STN-DBS treated patients, proprioceptive drift depended on baseline proprioceptive inaccuracy (the difference between the baseline estimate and the actual position of their middle finger); regression analysis showed an effect of stroking condition and an interaction between group and baseline proprioceptive inaccuracy (χ2 = 15.75, *p* < .001). Although baseline proprioceptive inaccuracy did not differ between controls and patients on- or off-stimulation (all *p* > .05), patients had less proprioceptive drift when their baseline estimates were closer to the position of rubber hand (interaction plot Fig. 3C). Examination of contrasts showed all subject groups had greater proprioceptive drift in the synchronous condition than in the asynchronous condition (χ2 = 44.91, *p* < .0001). Patients in the off-stimulation state had greater proprioceptive drift than controls in both stroking conditions (both χ2 = 7.95, *p* = .029). The difference between patients in the on-stimulation state and controls was not significant in either stroking condition (both *p* > .05). Contrary to our second hypothesis that switching STN-DBS on improves the precision of somatosensory perception and thus decreases proprioceptive drift, proprioceptive drift did not differ between on- and off-stimulation states in either stroking condition (both *p* > .05). Age, sex, cognition, mood and apathy were not significant covariates and thus excluded from the final model (Supplementary Table E1-2).

**Figure 3.**
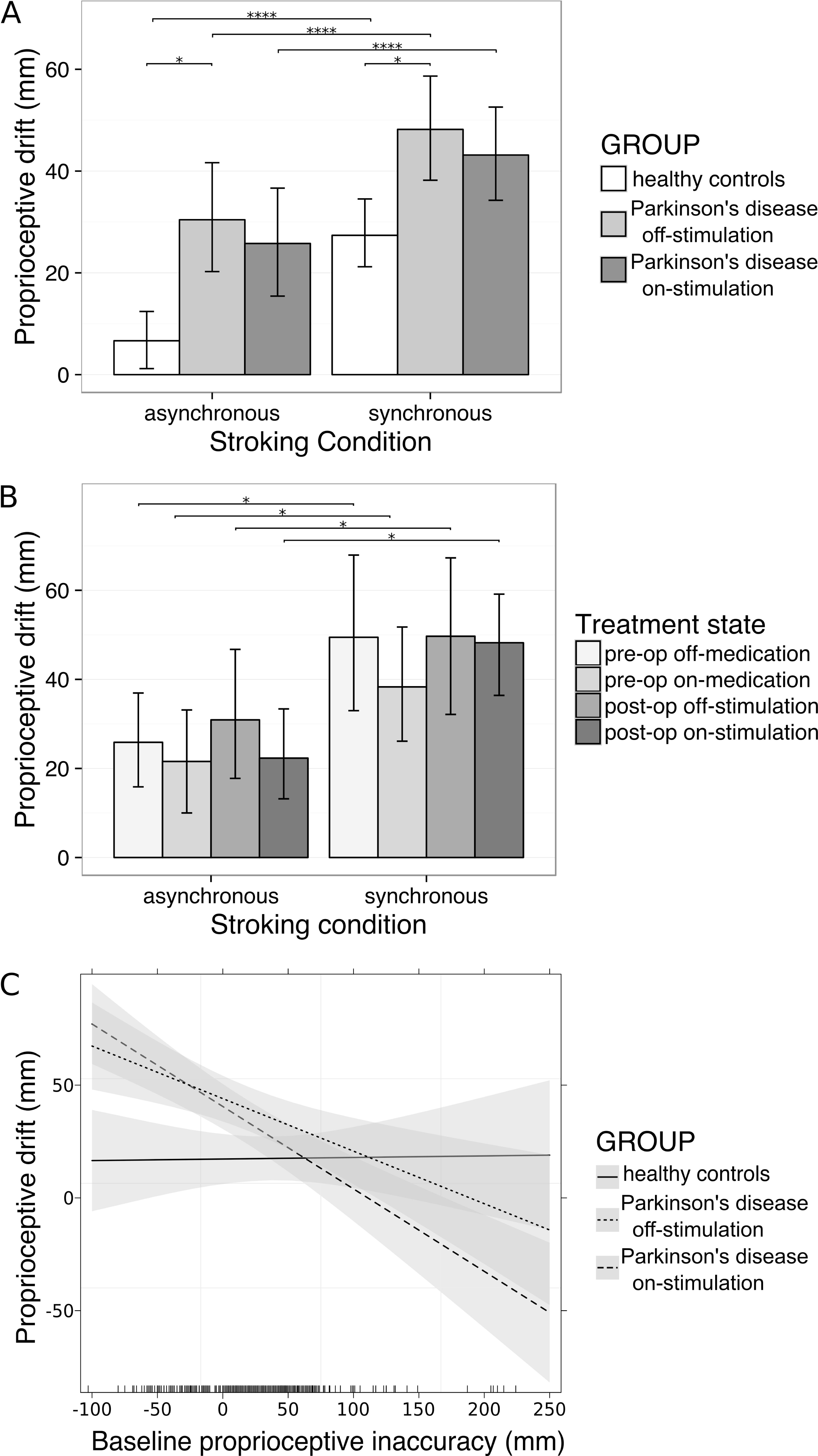
Mean and 95% CI of proprioceptive drift in A) 23 Parkinson’s disease patients treated with deep brain stimulation of the subthalamic nucleus (STN-DBS) and 21 controls, and B) a subset of 11 Parkinson's disease patients tested prior to STN-DBS implantation surgery (pre-op) whilst taking their usual anti-Parkinson drugs (‘on-medication’, n = 9), and after withholding all anti-Parkinson drugs (‘off-medication’, n = 11). Patients had greater proprioceptive drift than controls only in the off-stimulation state. Amongst the subset of patients studied preoperatively, there was no difference in proprioceptive drift between the four treatment states. P-values are based on the regression model, ****: p < .0001, * p <.05. All unlabelled contrasts were not significant. C) Interaction plot showing that in the 23 patients, proprioceptive drift (i.e. the difference between their baseline and post-stroking estimates of finger position) decreased as baseline estimates became more inaccurate towards the direction of the rubber hand. Shading: 95% CI.

#### Subset of PD patients studied preoperatively

The preoperative subset of 11 patients had similar results to the larger patient cohort (Fig. 3B). Proprioceptive drift was greater in the synchronous condition than in the asynchronous condition, and patients had less proprioceptive drift when their baseline estimates were closer to the position of rubber hand; regression analysis showed an effect of stroking condition (b = 20.93, 95% CI [0.53, 24.36], *p* = .016) and baseline proprioceptive inaccuracy (b = −0.17, 95% CI [−0.27, −0.07], *p* = .004). Examination of contrasts revealed no differences amongst the preoperative and postoperative treatments states (all *p* > .05). Duration of disease, motor severity and LEDD were not significant covariates and thus were not included in the final model (Supplementary Table F1-2).

### Reach movement metrics

Patients had a larger peak lateral displacement in the synchronous condition than in the asynchronous condition; regression analysis showed an effect of stroking condition (b = 11.32, 95% CI [3.60, 17.87], *p* = .004) and side-of-test (b = −45.91, 95% CI [−69.90, −13.08], *p* = .005) but not stimulation state (b = 0.59, 95% CI [-15.06, 15.80], *p* = .944). The effect of stroking condition was explained only by proprioceptive drift (b = 2.15, 95% CI [0.17, 4.31], *p* = .053), not post-stroking estimate (b = −0.47, 95% CI [-2.01, 1.21], *p* = .571) or motor severity (b = 0.03, 95% CI [-0.89, 1.07], *p* = .951). That is to say, when patients tried to touch the laterally positioned target, they made larger lateral movements in the synchronous condition than in the asynchronous condition because proprioceptive drift towards the medially placed rubber hand was greater in the synchronous condition.

No other movement metric was affected by stroking condition (Supplementary Table H-J). As expected, patients were faster in the on-stimulation state than in the off-stimulation state (mean velocity: b = 42.47, 95% CI [23.54, 69.22], *p* = .001; movement duration: b = −0.56, 95% CI [−0.91, −0.23], *p* = .006) The effects of stimulation on mean velocity and movement duration were explained by the improvement in motor severity (UPDRS) (Supplementary Table H-I).

## Discussion

PD and STN-DBS affect illusory perception in multisensory bodily awareness. Firstly, we replicated our previous findings—that people with PD do not reject the RHI as strongly as controls in the asynchronous condition, and have greater proprioceptive drift in both synchronous and asynchronous conditions—in a cohort of STN-DBS treated patients with more advanced disease. In the present study, we also modulated cortico-basal ganglia-thalamic circuitry by switching STN-DBS on and off. We found that switching STN-DBS on strengthens rejection of the RHI in the asynchronous condition, partially normalising patients’ experience of the illusion. By contrast, STN-DBS had no effect on proprioceptive drift. Surgical implantation of the STN-DBS electrodes alone, in the absence of electrical stimulation, had no effect on the RHI.

There are several possible explanations for why STN-DBS strengthens rejection of the RHI in the asynchronous condition. Rather than being all-or-nothing, the subjective RHI experience exists on a spectrum—it is strongest when temporal mismatch between visual and tactile cues is less than 300 ms, weakens with increasing mismatch, and is eliminated when mismatch is greater than 600 ms.^34^ Therefore, noisy or inaccurate temporal integration of visual and tactile cues in PD could lessen the asynchronous signal and weaken rejection of the RHI.

PD patients have well-characterised temporal deficits including increased somatosensory temporal detection thresholds and inaccurate estimation of temporal inter-stimulus intervals.^35,36^ Cope *et al.* (2014) found STN-DBS does not improve temporal discrimination of auditory inter-stimulus intervals in the hundreds of milliseconds range, however they only waited 5 min before testing their patients in the off-stimulation state.^37^ This may not be long enough for the effects of stimulation to wear off. Gulberti *et al.* (2015) found that chronic STN-DBS attenuated the effects of PD on auditory evoked potentials to rhythmic auditory stimuli, whereas transient interruptions to stimulation had no effect.^17^ Using an overnight stimulation “washout”, Koch *et al.* (2004) showed that PD patients discriminated auditory inter-stimulus intervals more accurately on-stimulation than off-stimulation.^11^ Extrapolating from these studies, STN-DBS may improve temporal discrimination in the RHI, strengthening the asynchronous signal, which would explain why our patients rejected the RHI more strongly in the asynchronous condition when in the ‘on-stimulation’ state.

However, improved temporal perception is insufficient to explain why STN-DBS strengthens rejection of the RHI in the asynchronous condition. Dopaminergic drugs also improve estimation of auditory inter-stimulus intervals^40^ but do not affect the RHI^5^. This may be because the RHI is more complex than auditory temporal estimation. Specifically, to arrive at a perceptual decision, the RHI requires not only integration of unimodal sensory signals with an internal representation of time, but also the integration of conflicting multisensory signals and higher order representations of body image.^1^ Notably, STN-DBS affected only the subjective questionnaire responses, but not the more implicit measures of proprioceptive drift and reach trajectory. This supports the theory that STN-DBS, but not dopaminergic drugs, alters complex cognitive^14,15^ and perceptual decisions.^16^ Simple cognitive and perceptual decisions are not affected by STN DBS. For example, Green *et al.* (2013) found that as visual stimulus complexity increased, the reaction times of STN-DBS treated patients did not increase as much as that of controls. This is also consistent with our study in (onset) binocular rivalry in which DBS, but not dopaminergic drugs, facilitated perceptual decisions.^38^ Moreover, this facilitation effect was seen in perceptual decisions during complex binocular rivalry, but not in simpler non-rivalrous situations.^38^

The STN is hypothesised to act as a brake in the cortical-striatal network, increasing the threshold of evidence required to make a complex decision when there are conflicting cues— a process thought to be disrupted by STN-DBS.^15^ How STN-DBS does this is not fully understood. Evidence from experiments in cognitive tasks points to modulation of brain network activity: EEG studies show that increased theta power in the medial prefrontal cortex associated with high-conflict tasks predicts increased time to decision, while STN-DBS reverses mediofrontal influence on decision thresholds.^17^ Furthermore, intraoperative intracranial recordings in the STN show corresponding changes in low-frequency power during high-conflict tasks, suggesting communication with the medial prefrontal cortex.^12^ In some cognitive and perceptual tasks, this can lead to increased errors,^14,16^ however in our study, switching stimulation on made perceptual inference closer to that of controls. There are similarities between our results and olfactory studies in PD, where STN-DBS ‘partially normalises’ odour recognition but not odour detection thresholds—suggesting that STN-DBS improves higher-order perceptual inference by modulating connectivity between the basal ganglia and sensory association cortices.^39,40^

The connection to the medial prefrontal cortex is part of the associative loop of the STN, anatomically and physiologically segregated from its motor and limbic loops.^6^ This associative loop may connect some of the neural substrates for multisensory integration in body awareness: fMRI activity in the ventromedial prefrontal cortex and lateral occipitotemporal cortex correlate with subjective RHI scores, whereas temporal synchronicity correlates with activation of the ventral premotor cortex.^41,42^ The ventral premotor cortex is also functionally connected to the basal ganglia in fMRI studies of temporal perception and prediction.^43^ These functional and anatomical connections may explain why STN-DBS strengthens rejection of the RHI in the asynchronous condition.

Consistent with our previous study in medically-treated PD patients,^5^ we found no evidence that PD or STN-DBS affects illusion scores in the illusion-promoting synchronous condition. We cannot exclude a ceiling effect on illusion scores in the present study or our previous one because the distribution of illusion scores in the synchronous condition skewed towards the upper limit of 14. The standard RHI questionnaire visual analogue scale may not capture the full spectrum of illusory experiences in PD patients.

We hypothesised that switching STN-DBS on reduces proprioceptive drift in both stroking conditions because stimulation has been reported to improve kinesthesia^9^ and haptic perception^10^ in PD. However, in our study, STN-DBS did not improve baseline or post-stroking estimates of finger position. Thus, any improvement in proprioceptive accuracy as a consequence of STN-DBS is probably insufficient to overcome the effects of visual dependence and illusory misperception.

Consistent with our final hypothesis that certain movement metrics would be affected by stroking condition, we found that when PD patients tried to touch the laterally positioned target, they made larger lateral movements in the synchronous condition than in the asynchronous condition. It is important to note that this was explained by proprioceptive drift (away from the reach target) rather than by their post-stroking estimate (i.e., how far away they perceived their hand to be from the target) or their motor severity (UPDRS). This finding establishes a connection between the atypical perceptual experience of the RHI in PD patients and bodily movement.

We found no effect of the RHI on the same movement metrics in our previous study in PD patients without STN-DBS^5^, but they may have been too mildly affected for such an effect to manifest. The STN-DBS treated patients in this study had longer disease durations, more motor severity (UPDRS off-treatment) and more proprioceptive drift. Proprioceptive deficits are known to impair movement in PD patients,^25^ and may make their movements more susceptible to somatic illusions. Our results are consistent with other studies that found somatic misperceptions influenced actions in PD. In these studies, sensory scaling errors reduced movement amplitude (hypokinesia)^44^, and teaching patients to make actions “too big” and vocalisations “too loud” overcame hypokinetic movement and speech that they perceive to be normal.^45^

Interpretation of our results should take into account the duration of stimulation “washout” in this study, which was 30 min. This was sufficient to invoke the effects that we report here and to elicit a significant change in UPDRS scores (Table 1), which is consistent with reports that 75% of the effect of STN-DBS on the signs of PD wears off within 15-30 minutes of discontinuation.^46^ Moreover, questionnaire responses and proprioceptive drift in the postoperative off-stimulation state (when chronic effects of stimulation might still be present) did not differ from preoperative values, which suggests that 30 min was sufficient for stimulation washout.

Despite postoperative confirmation that the DBS electrodes were situated in the STN, the precise location and active terminals were not uniform across patients, and we cannot predict individual differences in the spread of local electrical current. Hence we are unable to discern whether the effect of stimulation results from modulation of the STN or more wider cortico-basal ganglia-thalamic circuitry. Another effective target for DBS in PD is the GPi. In the future, it may be informative to compare the effects of DBS on the RHI in PD based on the exact site of stimulation (e.g., STN vs GPi).

In conclusion, illusory misperception of arm position was greater in patients with severe PD than healthy subjects, and altered their subsequent movements. Stimulation of the STN had no effect on proprioceptive drift. However, not only have we confirmed that PD patients are less certain about rejecting illusory experience than healthy subjects in a typically unambiguous condition, we discovered that STN-DBS partially normalises their responses. Our findings implicate the STN and subcortical connections in the network for multisensory integration in bodily awareness and pave the way for future research into the perceptual effects of DBS.

## Acknowledgements

The authors wish to thank Richard Beare for help with R statistical software.

## Author Contributions Statement

All authors contributed to the design of the project. C.D wrote the first draft and prepared the figures. All authors reviewed and edited the manuscript.

## References

1. Ehrsson, H. H. The concept of body ownership and its relation to multi-sensory integration. In The new handbook of multisensory processing. 775 – 792 (MIT Press., 2012).

2. Ehrsson, H. H., Spence, C. & Passingham, R. E. That’s my hand! Activity in premotor cortex reflects feeling of ownership of a limb. Science 305, 875–877 (2004).

3. Ehrsson, H. H., Holmes, N. P. & Passingham, R. E. Touching a rubber hand: feeling of body ownership is associated with activity in multisensory brain areas. J. Neurosci. 25, 10564–10573 (2005).

4. Kammers, M. P. M., de Vignemont, F., Verhagen, L. & Dijkerman, H. C. The rubber hand illusion in action. Neuropsychologia 47, 204–211 (2009).

5. Ding, C. et al. Parkinson’s disease alters multisensory perception: Insights from the Rubber Hand Illusion. Neuropsychologia 97, 38–45 (2017).

6. DeLong, M. R. & Wichmann, T. Circuits and circuit disorders of the basal ganglia. Arch. Neurol. 64, 20–24 (2007).

7. Herrington, T. M., Cheng, J. J. & Eskandar, E. N. Mechanisms of deep brain stimulation. J. Neurophysiol. 115, 19–38 (2016).

8. Spielberger, S., Wolf, E., Kress, M., Seppi, K. & Poewe, W. The influence of deep brain stimulation on pain perception in Parkinson’s disease. Mov. Disord. 26, 1367–8; author reply 1368-9 (2011).

9. Maschke, M., Tuite, P. J., Pickett, K., Wächter, T. & Konczak, J. The effect of subthalamic nucleus stimulation on kinaesthesia in Parkinson’s disease. J. Neurol. Neurosurg. Psychiatry 76, 569–571 (2005).

10. Aman, J. E., Abosch, A., Bebler, M., Lu, C.-H. & Konczak, J. Subthalamic nucleus deep brain stimulation improves somatosensory function in Parkinson’s disease. Mov. Disord. 29, 221–228 (2014).

11. Koch, G. et al. Subthalamic deep brain stimulation improves time perception in Parkinson’s disease. Neuroreport 15, 1071–1073 (2004).

12. Gulberti, A. et al. Predictive timing functions of cortical beta oscillations are impaired in Parkinson’s disease and influenced by L-DOPA and deep brain stimulation of the subthalamic nucleus. Neuroimage Clin 9, 436–449 (2015).

13. Halpern, C. H., Rick, J. H., Danish, S. F., Grossman, M. & Baltuch, G. H. Cognition following bilateral deep brain stimulation surgery of the subthalamic nucleus for Parkinson’s disease. Int. J. Geriatr. Psychiatry 24, 443–451 (2009).

14. Weintraub, D. B. & Zaghloul, K. A. The role of the subthalamic nucleus in cognition. Rev. Neurosci. 24, 125–138 (2013).

15. Frank, M. J., Samanta, J., Moustafa, A. A. & Sherman, S. J. Hold your horses: impulsivity, deep brain stimulation, and medication in parkinsonism. Science 318, 1309–1312 (2007).

16. Green, N. et al. Reduction of influence of task difficulty on perceptual decision making by STN deep brain stimulation. Curr. Biol. 23, 1681–1684 (2013).

17. Gulberti, A. et al. Subthalamic deep brain stimulation improves auditory sensory gating deficit in Parkinson’s disease. Clin. Neurophysiol. 126, 565–574 (2015).

18. Schmalbach, B. et al. The subthalamic nucleus influences visuospatial attention in humans. J. Cogn. Neurosci. 26, 543–550 (2014).

19. Witt, K., Kopper, F., Deuschl, G. & Krack, P. Subthalamic nucleus influences spatial orientation in extra-personal space. Mov. Disord. 21, 354–361 (2006).

20. Kammers, M. P. M., Kootker, J. A., Hogendoorn, H. & Dijkerman, H. C. How many motoric body representations can we grasp? Exp. Brain Res. 202, 203–212 (2010).

21. Heed, T. et al. Visual information and rubber hand embodiment differentially affect reach-to-grasp actions. Acta Psychol. 138, 263–271 (2011).

22. Palmer, C. J., Paton, B., Hohwy, J. & Enticott, P. G. Movement under uncertainty: the effects of the rubber-hand illusion vary along the nonclinical autism spectrum. Neuropsychologia 51, 1942–1951 (2013).

23. Zopf, R., Truong, S., Finkbeiner, M., Friedman, J. & Williams, M. A. Viewing and feeling touch modulates hand position for reaching. Neuropsychologia 49, 1287–1293 (2011).

24. Mongeon, D., Blanchet, P., Bergeron, S. & Messier, J. Impact of Parkinson’s disease on proprioceptively based on-line movement control. Exp. Brain Res. 233, 2707–2721 (2015).

25. Lee, D., Henriques, D. Y., Snider, J., Song, D. & Poizner, H. Reaching to proprioceptively defined targets in Parkinson’s disease: effects of deep brain stimulation therapy. Neuroscience 244, 99–112 (2013).

26. Palmer, C. J., Paton, B., Kirkovski, M., Enticott, P. G. & Hohwy, J. Context sensitivity in action decreases along the autism spectrum: a predictive processing perspective. Proc. Biol. Sci. 282, (2015).

27. (R Core Team). R: A language and environment for statistical computing. (2014). Available at: http://www.R-project.org/. (Accessed: 19th July 2016)

28. Bates, D., Maechler, M., Bolker, B. & Walker, S. lme4: Linear mixed-effects models using Eigen and S4. R package version 1.1-7. (2014). Available at: http://CRAN.R-project.org/package=lme4. (Accessed: 19th July 2016)

29. Pinheiro, J., Bates, D., DebRoy, S., Sarkar, D. & (R Core Team). nlme: Linear and Nonlinear Mixed Effects Models. R package version 3.1-117. (2014). Available at: http://CRAN.R-project.org/package=nlme. (Accessed: 19th July 2016)

30. Fox, J. & Hong, J. Effects: Effect Displays in R for Multinomial and Proportional-Odds Logit Models: Extensions to the effects Package. Journal of Statistical Software, 32(1), 1–24. (2009). Available at: http://www.jstatsoft.org/v32/i01/. (Accessed: 19th July 2016)

31. Fox, J. Effects: Effect Displays in R for Generalised Linear Models. Journal of Statistical Software, 8(15), 1–27. (2003). Available at: http://www.jstatsoft.org/v08/i15/. (Accessed: 19th July 2016)

32. De Rosario-Martinez, H. phia: Post-Hoc Interaction Analysis R package version 0.2-1. (2015). Available at: http://CRAN.R-project.org/package=phia. (Accessed: 19th July 2016)

33. Hlavac, M. stargazer. stargazer: Well-Formatted Regression and Summary Statistics Tables. R package version 5.2 (2015). Available at: http://CRAN.R-project.org/package=phia.

34. Shimada, S., Fukuda, K. & Hiraki, K. Rubber hand illusion under delayed visual feedback. PLoS One 4, e6185 (2009).

35. O’Boyle, D. J., Freeman, J. S. & Cody, F. W. The accuracy and precision of timing of self-paced, repetitive movements in subjects with Parkinson’s disease. Brain 119 (Pt 1), 51–70 (1996).

36. Pastor, M. A., Artieda, J., Jahanshahi, M. & Obeso, J. A. Time estimation and reproduction is abnormal in Parkinson’s disease. Brain 115 Pt 1, 211–225 (1992).

37. Cope, T. E. et al. Subthalamic deep brain stimulation in Parkinson's disease has no significant effect on perceptual timing in the hundreds of milliseconds range. Neuropsychologia 57, 29–37 (2014).

38. Fujiwara, M. et al. Optokinetic nystagmus reflects perceptual directions in the onset binocular rivalry in Parkinson’s disease. PLoS One 12, e0173707 (2017).

39. Guo, X. et al. Effects of bilateral deep brain stimulation of the subthalamic nucleus on olfactory function in Parkinson’s disease patients. Stereotact. Funct. Neurosurg. 86, 237–244 (2008).

40. Hummel, T., Jahnke, U., Sommer, U., Reichmann, H. & Müller, A. Olfactory function in patients with idiopathic Parkinson’s disease: effects of deep brain stimulation in the subthalamic nucleus. J. Neural Transm. 112, 669–676 (2005).

41. Bekrater-Bodmann, R. et al. The importance of synchrony and temporal order of visual and tactile input for illusory limb ownership experiences - an FMRI study applying virtual reality. PLoS One 9, e87013 (2014).

42. Limanowski, J., Jakub, L. & Felix, B. That’s not quite me: limb ownership encoding in the brain. Soc. Cogn. Affect. Neurosci. nsv079 (2015).

43. Grahn, J. A. & Rowe, J. B. Feeling the beat: premotor and striatal interactions in musicians and nonmusicians during beat perception. J. Neurosci. 29, 7540–7548 (2009).

44. Berardelli, A., Rothwell, J. C., Thompson, P. D. & Hallett, M. Pathophysiology of bradykinesia in Parkinson’s disease. Brain 124, 2131–2146 (2001).

45. Fox, C., Ebersbach, G., Ramig, L. & Sapir, S. LSVT LOUD and LSVT BIG: Behavioral Treatment Programs for Speech and Body Movement in Parkinson Disease. Parkinsons Dis. 2012, 391–946 (2012).

46. Temperli, P. et al. How do parkinsonian signs return after discontinuation of subthalamic DBS? Neurology 60, 78–81 (2003).

